# Scalable projected Light Sheet Microscopy for high-resolution imaging of living and cleared samples

**DOI:** 10.1101/2023.05.31.543173

**Authors:** Yannan Chen, Shradha Chauhan, Cheng Gong, Hannah Dayton, Cong Xu, Estanislao Daniel De La Cruz, Malika S. Datta, Kam W. Leong, Lars E.P. Dietrich, Raju Tomer

## Abstract

Light sheet fluorescence microscopy (LSFM) is a widely used imaging technique for living and large cleared samples. However, high-performance LSFM systems are often prohibitively expensive and not easily scalable for high-throughput applications. Here, we introduce a cost-effective, scalable, and versatile high-resolution imaging framework, called projected Light Sheet Microscopy (pLSM), which repurposes readily available off-the-shelf consumer-grade components and an over-the-network control architecture to achieve high-resolution imaging of living and cleared samples. We extensively characterize the pLSM framework and showcase its capabilities through high-resolution, multi-color imaging and quantitative analysis of mouse and post-mortem human brain samples cleared using various techniques. Moreover, we show the applicability of pLSM for high-throughput molecular phenotyping of human induced pluripotent cells (iPSC)-derived brain and vessel organoids. Additionally, we utilized pLSM for comprehensive live imaging of bacterial pellicle biofilms at the air-liquid interface, uncovering their intricate layered architecture and diverse cellular dynamics across different depths. Overall, the pLSM framework has the potential to further democratize LSFM by making high-resolution light sheet microscopy more accessible and scalable.

## INTRODUCTION

Light sheet fluorescence microscopy (LSFM) has emerged as a powerful and versatile approach for biological imaging, enabling prolonged *in vivo* observations and facilitating high-resolution imaging of large cleared intact organs^1^. These remarkable capabilities are achieved through LSFM’s distinctive orthogonal planar illumination and detection geometry, which minimizes energy load and allows use of fast CMOS camera-based wide-field detection. In recent years, several highly optimized LSFM implementations have been developed for high-speed phenotyping of intact cleared organs at high resolution^1–10^ and high-speed live imaging of cells, embryos and functioning nervous systems^3, 11–15^. However, the substantial costs associated with implementing these systems and their limited scalability hinder their utility in high-throughput imaging applications and impede broader accessibility within the scientific community. To address some of these constraints, several open-source initiatives have emerged, including openSPIM^16^, OpenSpinMicroscopy^17^ and mesoSPIM^18^, which aim to optimize the cost and portability of LSFM. Nevertheless, there is still an unmet need for further cost-optimized implementations and genuinely scalable systems that can operate remotely over networks to facilitate high-throughput applications such as drug screening and the characterization of diseased brains.

In this study, we introduce a versatile LSFM imaging framework called projected LSFM (pLSM), which offers a cost-effective and scalable solution without compromising imaging quality. Our approach leverages off-the-shelf components, including pocket LASER projectors as multi-color illumination sources, Nvidia Jetson Nano boards for electronic control, 3D-printed imaging chambers, and optimized scan and detection optics. Additionally, we have developed a network-based imaging workflow using native Python and Jupyter Notebook, enabling remote control of multiple microscopes at scale.

In our extensive characterization of the pLSM system, we demonstrate its robustness and versatility by performing high-resolution multi-color imaging of large mouse and human brain samples cleared using various methods, enabling whole-brain quantitative phenotyping comparable to high-end LSFM systems. Moreover, we establish high-throughput pLSM imaging capabilities for a large number of human iPSC-derived brain and vessel organoids at sub-cellular resolution, addressing their sample-to-sample variabilities. Furthermore, we highlight the unique advantages of pLSM for live imaging of bacterial cellular dynamics in pellicle biofilms at the air-liquid interface, revealing crucial insights into their dynamic layered architecture.

Overall, our study highlights the versatility, cost-effectiveness, and high-quality imaging capabilities of the pLSM framework in various applications, ranging from brain imaging to bacterial biofilms at air-liquid interface.

## RESULTS

### pLSM implementation

A LSFM system encompasses orthogonal illumination and detection optics, laser light sources, control electronics, and data acquisition software^1^. Our goal was to optimize the cost-effectiveness of each component by incorporating readily available, low-cost consumer-grade parts, while ensuring high imaging quality. The pLSM system comprises two identical modules for light-sheet illumination, which collectively provide comprehensive coverage of large cleared samples, along with an orthogonal wide-field detection arm for image detection (**Fig. 1a-b, Supplementary Table 1**). Additionally, 3D motorized stages are employed for multi-tile imaging of large samples mounted in quartz glass cuvettes^2, 3^ within a 3D-printed immersion chamber. To implement the electronic controls and imaging workflow, we employed a cost-effective Nvidia Jetson Nano board, which can be operated remotely via an SSH-based connection and a Python-based control software GUI within the Jupyter Notebook framework (**Fig. 1c**).

**Figure 1.**
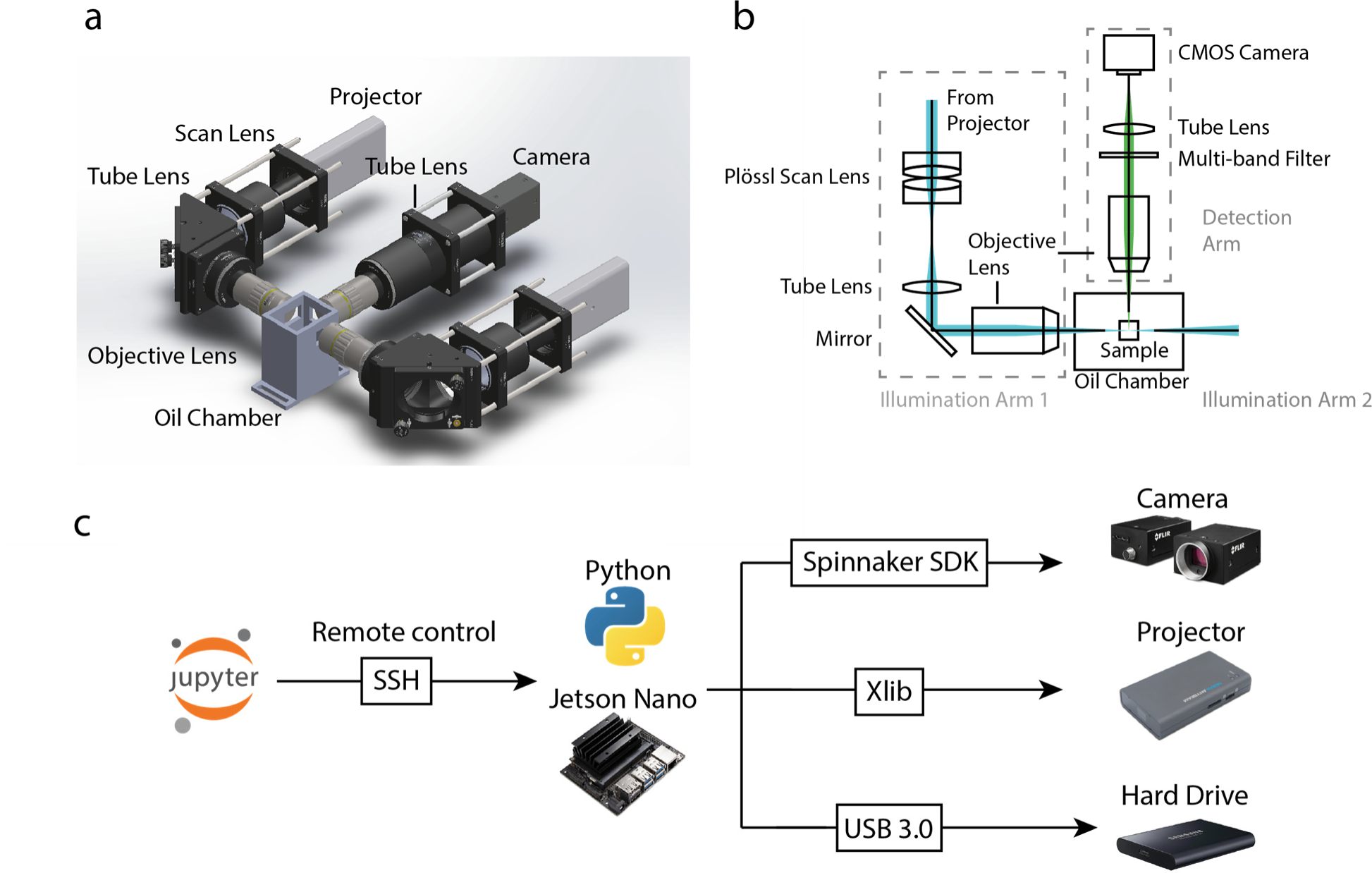
Implementation of projected Light Sheet Microscopy. (a-b) 3D model and optical paths of the pLSM system. The pLSM system repurposes pocket LASER projectors for generation of thin scanned light sheets of up to three wavelengths. Two opposing light sheets are employed to illuminate the sample, which is positioned within an oil chamber (the motorized stage holding the sample cuvette is not depicted). The illuminated selective plane is captured by an orthogonally arranged in-focus detection objective, tube lens, and a consumer-grade CMOS camera. (c) The control code, developed using Python, is executed on an Nvidia Jetson Nano board. Remote operation of the system is enabled through a Jupyter Notebook-based graphical user interface (GUI) utilizing the SSH remote protocol. The FLIR CMOS camera is controlled by the Spinnaker SDK package, while the LASER projector via the Xlib library to generate the scanned light sheets. The acquired camera data is streamed to a solid-state hard drive through a high-speed USB 3.0 port.

For the detection arm, we utilized a long-working distance air objective (Mitutoyo 10x Plan Apo) or multi-immersion objective (ASI 16.67X/0.4NA), a 200 mm focal length tube lens (TTL-200, Thorlabs), a multi-band pass emission filter, and a low-cost consumer-grade CMOS camera (GS3-U3-89S6M-C, FLIR, 3.45 µm pixel) for wide-field detection of the illuminated plane. Notably, this camera provided performance comparable to high-end sCMOS camera, albeit with lower quantum efficiency. Next, we sought to develop cost-effective alternatives to expensive multi-laser engines, galvo scanners, and associated controllers for generation of thin illumination light sheets. To achieve this, we devised an approach that employs pocket laser projectors for generation of high-quality scanned light sheets with 3-color imaging capabilities (**Fig. 1**). We found that these economical projectors provide substantial benefits of fully integrated 3-color laser diodes (TO-8 set with blue: 440-460 nm, green: 515-530 nm, and red: 632-642 nm) with sufficient power needed for imaging, MEMS 2D-scanners, and associated electronics in a compact assembly. However, adapting them for light-sheet generation presented practical challenges in controlling the various light-sheet properties, positioning, and scanning speeds. To address these challenges, we devised a control framework utilizing the Python X library (Python-Xlib) to directly modulate various scanning parameters (**Fig. 1c**). This framework allows the generation of light sheets as multi-pixel (1 or more, on-demand) wide projected lines of specific height and location (**Fig. 2a-d**). Importantly, this approach also offers the unique advantage of easily modulating the light sheet’s thickness and field-of-view (FOV) by controlling the pixel widths of the projected line (**Fig. 2d**). Downstream to the projector, we employed a Plössl lens (implemented as a pair of achromats, AC254-045-A) as an effective substitute for expensive f-theta scan lenses, an achromat doublet (AC508-075-A) as a tube lens, and an air illumination objective (Mitutoyo 10x Plan Apo).

**Figure 2.**
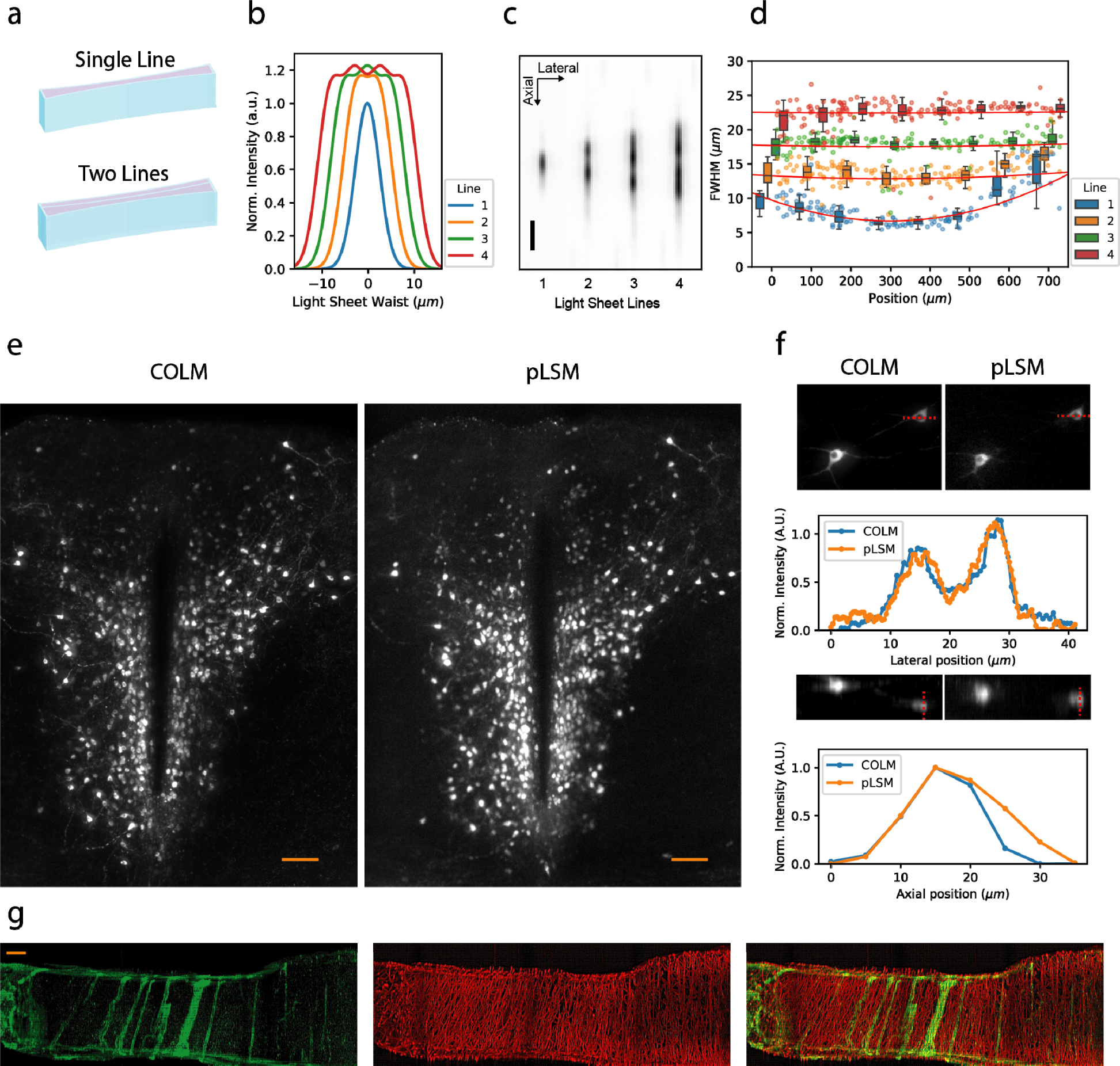
Characterization of projected Light Sheet Microscopy. (a) Schematic illustration demonstrating the generation of light sheets by projecting lines of varying pixel widths (shown for 1- and 2-pixels wide lines). (b) Simulated intensity profiles of the light sheet waist generated by projection of different pixel-width lines. Notably, increasing the number of lines improves the uniformity of the light sheet throughout its cross-section. (c) Measurement of the illumination Point Spread Function (PSF) for the light sheet generated by varying pixel widths. The scale bar represents 10µm. (d) The PSF measurements across the field of view (FOV). In this implementation of pLSM, a single-pixel wide light sheet achieves a thickness of approximately 5 µm at the waist. Light sheets with multiple pixels exhibit greater thickness while achieving significantly improved uniformity across the FOV. (e) Maximum Intensity Projection (MIP) images of TH+ neurons obtained from the same sample using pLSM (with a 2-pixel wide light sheet) and high-end LSFM implementation COLM. Scale bar represents 100µm. (f) Quantitative analysis and comparison of the intensity profiles of a cell imaged by pLSM and COLM. The pLSM image was preprocessed by applying median filtering (radius=0.1) followed by Gaussian filtering (σ=1) to eliminate hot pixels. (g) Multicolor imaging of human cortex vasculature, stained with Acta2 (AF555, Green) and Podocalyxin/CD31 (AF647, Red). The scale bar represents 200µm.

The pLSM control framework and imaging workflows were implemented on a cost-effective Nvidia Jetson Nano board running the Linux operating system. The Spinnaker SDK package for image grabbing and Python-Xlib for light sheet projection. The acquired images are streamed to a USB 3.0 high-speed solid-state hard drive. The control GUI was implemented using IPyWidgets within the Jupyter Notebook framework, enabling remote operation via a SSH connection from any machine. This network-controlled framework allows for easy scalability without requiring the full capacity of a remote machine.

### pLSM characterization

We first performed a comprehensive characterization of the light sheets generated through the utilization of multi-pixel projected lines in our pLSM framework. We imaged micron-size fluorescent beads to characterize the light sheet and detection point spread functions. As shown in **Fig. 2c-d**, in this implementation of the pLSM, the generated light sheets have a thickness of ∼5 µm when using a single pixel projected line. Furthermore, for larger pixel widths projected lines, we observed sub-linear increases in the thickness of the light sheet (**Fig. 2d**). Importantly, the modulation of light sheet thickness and field-of-view can be easily accomplished at the software level, offering a convenient means for rapid adaptation to diverse experimental requirements. Additionally, our findings indicate that this approach yields a flatter intensity profile across the light sheet cross-section compared to a standard Gaussian light sheet of similar thickness (**Fig. 2b-d**). Notably, the scanning speed of the light sheet, and consequently the imaging speed, is solely determined by the projector refresh rate (60 Hz for the utilized model), independent of the pixel widths employed. As a result, pLSM presents a highly adaptable multi-scale scheme capable of accommodating various imaging needs, spanning from high axial resolution settings to high-volume acquisition rates with thicker light sheets.

To further evaluate the imaging performance of pLSM, we conducted a direct comparison with a high-end COLM^2, 3^ system. To this end, we utilized the same cleared mouse brain sample stained for dopaminergic neurons and acquired images using both pLSM and COLM implementations. Quantification of the intensity distribution, as presented in **Fig 2e-f**, demonstrates that pLSM achieves imaging quality comparable to that of the high-end COLM implementation. This result highlights the capabilities of pLSM in delivering high-resolution imaging outcomes on par with established state-of-the-art methodologies. Furthermore, we demonstrate the multi-color imaging capability of pLSM by performing 2-channel imaging of a post-mortem human brain sample stained with two different markers of vasculatures (**Fig. 2g**) by using green (515-530 nm) and red (632-642 nm) light. Overall, pLSM allows high-resolution imaging of biological samples with comparable quality to high-end implementations.

In summary, the detailed characterization demonstrates that pLSM provides high-quality imaging comparable to high-end and expensive LSFM implementations, while offering a flexible, cost-effective, and scalable platform for high-resolution imaging of biological samples.

### High-resolution mapping of large intact samples cleared with multiple clearing techniques

To enable high quality imaging of large intact samples, we optimized the imaging workflow of pLSM to address the sample-induced optical aberrations at different locations. A key improvement was the implementation of a linearly adapting light-sheet offset correction procedure. By sampling optimal parameters at a few (typically 5-10) anchor positions and applying 3D linear interpolation, the pLSM system generates a smoothed light-sheet offsets map to compensate for optical shifts resulting from refractive index mismatches. Importantly, this approach ensures that pLSM remains compatible with various tissue clearing methods.

We conducted high-resolution imaging of intact mouse brain samples cleared using the iDISCO^19^ and CLARITY^2, 20^ techniques. In one experiment, our focus was on mapping the dopamine (DA) system throughout the entire mouse brain using pLSM imaging. Intact mouse brain samples were stained with an antibody against tyrosine hydroxylase (TH), a widely used marker of DA neurons, and an Alexa Fluor 647 secondary antibody. The samples were cleared using the iDISCO method and imaged with red projected lines (632-642 nm) and a Mitutoyo 10x (Plan Apo) objective. The results (**Fig. 3a** and **Supplementary videos 1**) demonstrate successful high-quality imaging of the intact mouse brain sample, allowing for the complete reconstruction of the DA system. Additionally, we utilized pLSM (with same specifications) to map the entire micro-vasculature in an intact mouse brain (**Fig. 3b**). The vasculature labeling was achieved by utilizing a cocktail of antibodies against Podocalyxin, CD31, and Acta2 (all detected using the same Alexa 647 secondary antibody)^21^.

**Figure 3.**
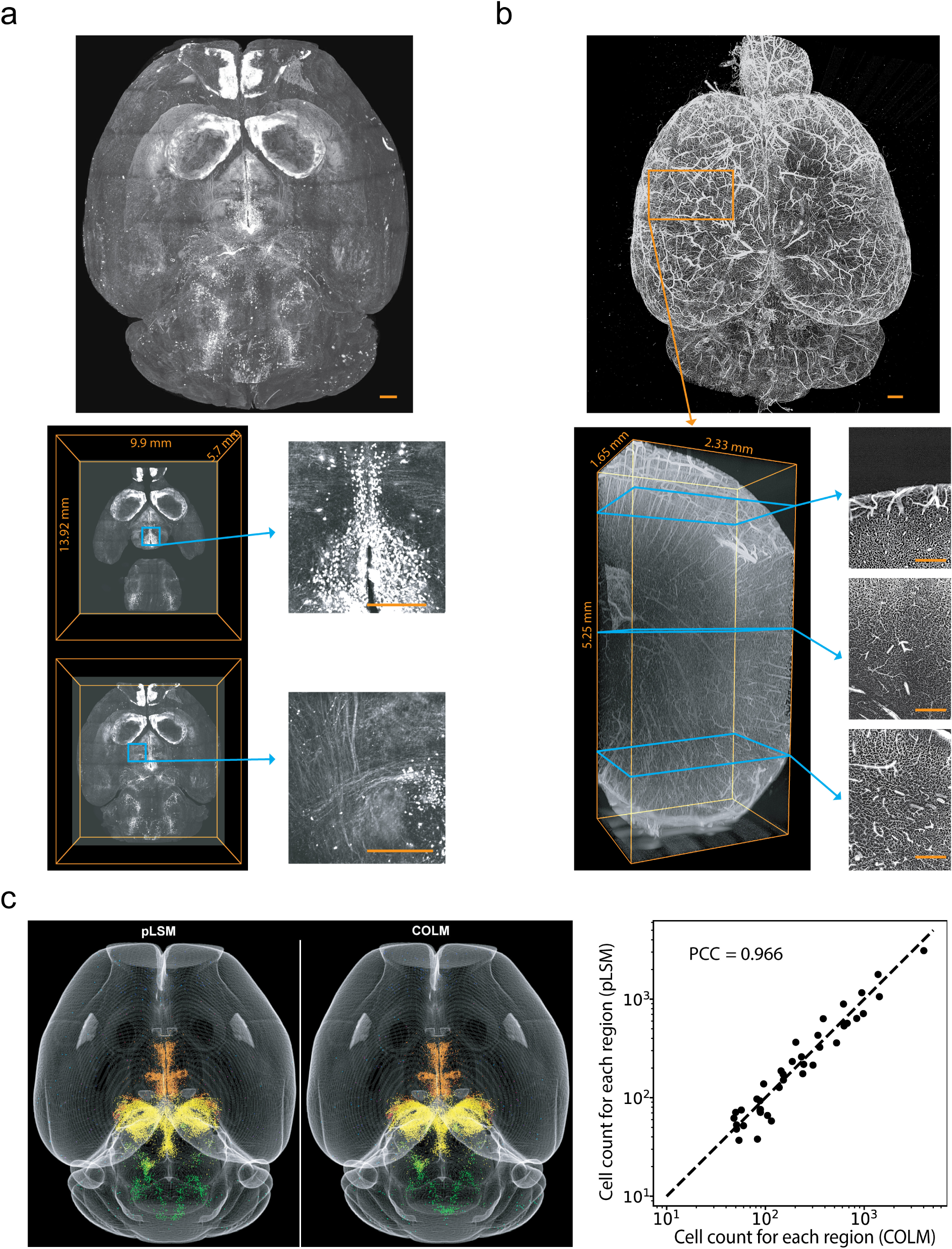
High-resolution pLSM imaging and quantitative analysis of intact mouse brain samples cleared with iDISCO. (a) High-resolution imaging of TH+ dopamine neurons in an iDISCO+ cleared brain sample. The top image presents an overview of the entire brain, while the bottom images provide visualizations of specific neuronal features at various depths. Scale bars represent 500 µm. Also see **Supplementary Videos 1-2** (b) High-resolution imaging of the complete mouse brain vasculature in an iDISCO cleared sample. The top image illustrates the rendering of the entire brain, while the bottom-left image exhibits a 3D rendering of a smaller volume. Additionally, 100µm thickness Maximum Intensity Projections (MIPs) are depicted at different imaging depths. Scale bars represent 500 µm. (c) Segmentation of TH+ dopamine neurons in datasets acquired by pLSM and COLM imaging of the same sample. The point clouds are color-coded according to the regional annotations from the Allen Brain Atlas at level 5. The right figure shows comparative analysis of region-specific neuron counts extracted from pLSM and COLM images. The top 40 regions at ABA level 8 were included in the analysis. The Pearson correlation coefficient of 0.966 indicates a high degree of agreement between the two imaging methods.

We further investigated whether the whole brain datasets acquired using pLSM could provide quantitative insights comparable to those obtained from high-end LSFM implementations. To explore this, we performed high-resolution imaging of the same TH-labeled mouse brain sample using both pLSM and the COLM^2^ system (**Fig. 3c, Supplementary Video 2**). We utilized the suiteWB pipeline^22^ to map TH+ DA neurons. The point clouds generated from the two systems showed strong agreements across all brain regions (**Fig. 3c**), demonstrating the compatibility of pLSM-acquired data with whole brain analysis approaches.

Lastly, to assess the compatibility of pLSM with different tissue clearing methods, we performed whole-brain imaging of CLARITY-cleared mouse brain samples from the *Thy1-eYFP* transgenic mouse line. The samples were cleared using the passive CLARITY method^2^ and imaged with green-color (515-530 nm) projected lines and Mitutoyo 10x objective. As shown in **Fig. 4a** and **Supplementary Video 3**, pLSM achieved high-resolution imaging of CLARITY cleared intact mouse brain. Additionally, we demonstrated the compatibility of pLSM with different detection objectives by performing high-resolution imaging with the ASI 16.67X/0.4 NA objective (**Fig. 4b, Supplementary Video 4**).

**Figure 4.**
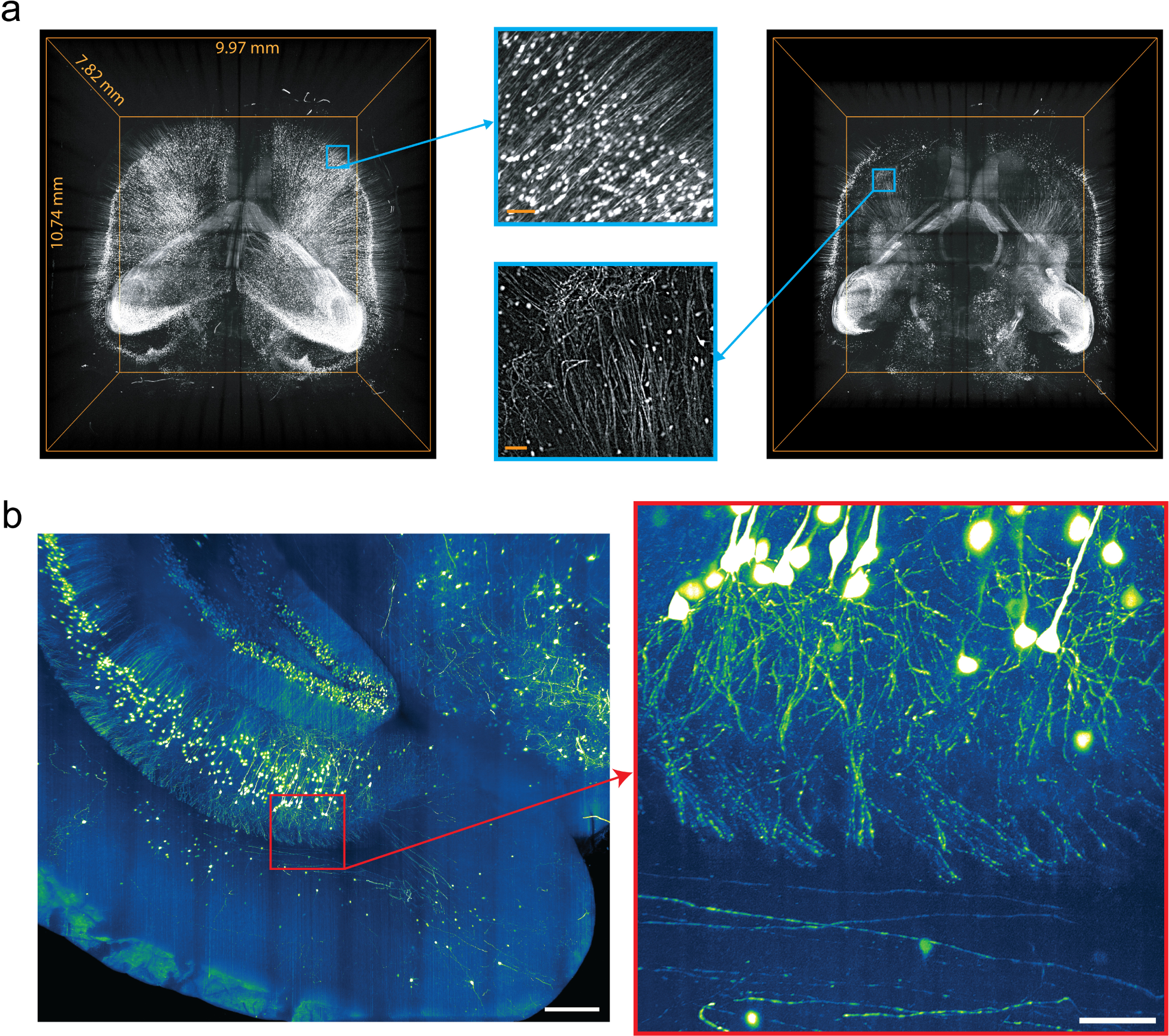
High-resolution pLSM imaging of CLARITY cleared mouse *Thy1-eYFP* brain. (a) High-resolution imaging of an intact *Thy1-eYFP* mouse brain cleared through passive CLARITY technique. The left image presents the rendering of the whole brain images, and the right image shows a 3D rendering of the deeper half of the brain. Also see **Supplementary Video 3**. Scale bars represent 100 µm. (b) High-resolution imaging of CLARITY cleared *Thy1-eYFP* mouse brain sample with ASI 16.67X/0.4 NA objective. Also see **Supplementary Video 4**. Scale bars represent 500 µm (left) and 100 µm (right).

In conclusion, these imaging experiments substantiate the capability of pLSM for high-resolution imaging of large intact samples cleared using multiple methods. The incorporation of the linearly adapting light-sheet offset correction procedure effectively compensated for the misalignments of detection focal plane with the illumination light sheet, ensuring high quality imaging throughout the sample. Notably, despite its cost-effective nature, pLSM is capable of generating quantitative insights comparable to those obtained from high-end LSFM systems.

### High-throughput phenotyping of human iPSC-derived brain and vessel organoids

Recent advances in brain and vessel organoids are enabling unprecedented access to some of the molecular, cellular and developmental mechanisms underlying complex diseases^23, 24^. However, these *in vitro* 3D preparations often exhibit considerable sample-to-sample variability, necessitating the use of large sample sizes for robust comparisons across different experimental conditions. In this study, we demonstrate the effectiveness of the scalable pLSM framework for high-throughput imaging of a large number of organoid samples at sub-cellular resolution (**Fig. 5, Supplementary Video 5**).

**Figure 5.**
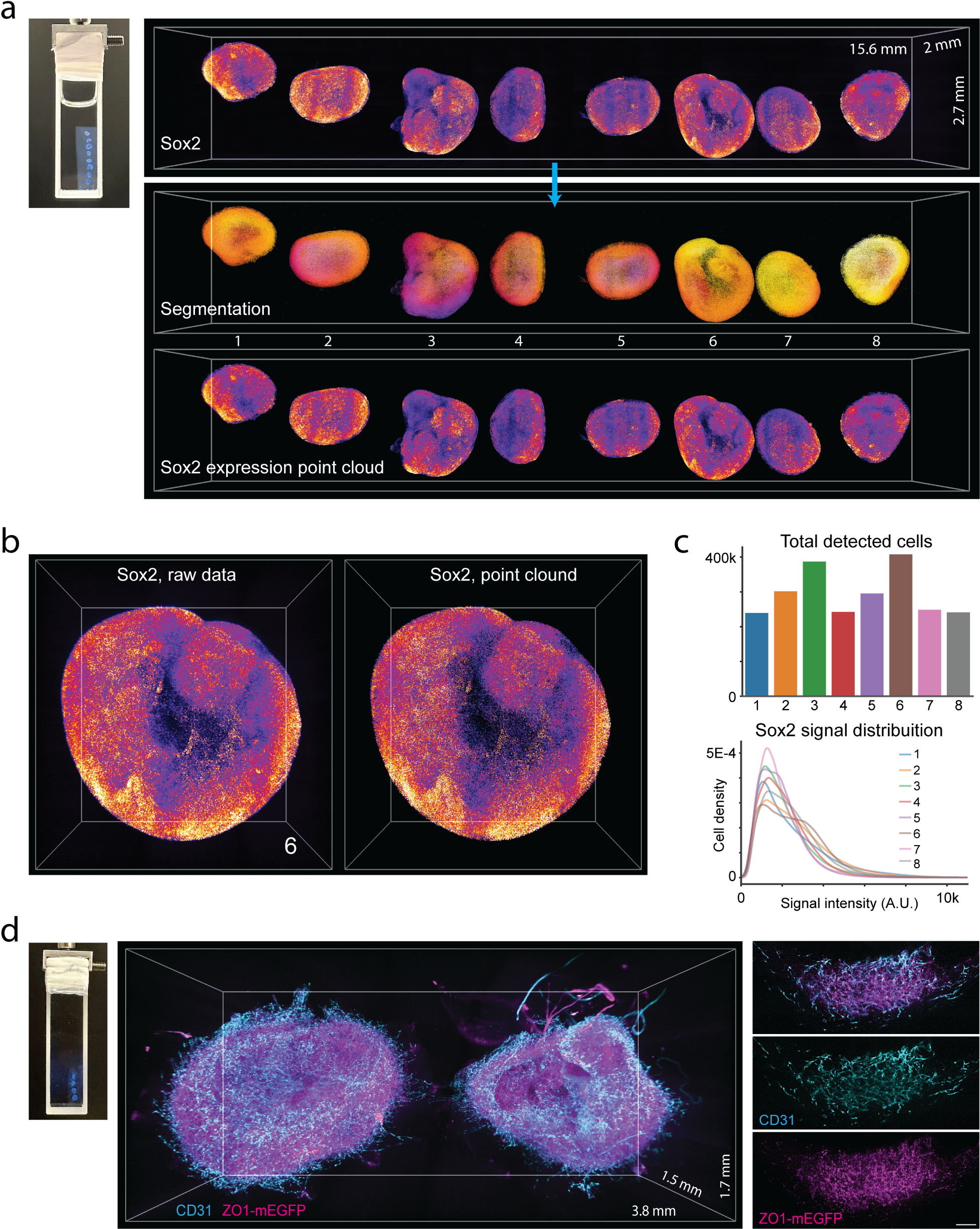
High-throughput phenotyping of human iPSC-derived brain and vessel organoids. (a) Mounting and pLSM imaging of an ensemble of brain organoid samples, immunostained with anti-Sox2 antibody and cleared with FDISCO technique. The volumetric renderings show the raw data (top), segmentation (middle) and Sox2 expression point clouds (bottom). The bounding box is 15.6 x 2 x 2.7 mm^3^. Also see **Supplementary Video 5**. (b) A zoomed-in view of brain organoid sample #6, showing raw data and the reconstructed expression point cloud. The bounding box is 1.84 x 1.935 x 1.13 mm^3^. (c) Quantification of total number of detected cells and Sox2 signal intensity distribution, showing the variability across brain organoid samples generated in same batch. (d) Mounting and multi-color pLSM imaging of vessel organoids stained with anti-CD31 antibody and ZO1-mEGFP reporter expression. The peripheral sprouting vessels exhibit lower ZO1-mEGFP signal. The scale bar is 100 µm. Also see **Supplementary Video 6**.

We first optimized the sample mounting for rapid imaging of a large ensemble of organoids in a single imaging session (**Fig. 5a, Supplementary Video 5**). This was achieved by immobilizing labelled organoid samples in a line by embedding them in 1% agarose gel^25^, which can then be processed with various tissue clearing methods, including the FDISCO^26^ method used in these experiments. The cleared organoids ensemble was mounted in a cuvette for unhindered optical access from all sides and imaged with pLSM at high resolution. This approach allowed us to perform high-throughput mapping of Sox2 (a marker of progenitor cells) expression in 8 brain organoid samples generated in a single batch. As shown in **Fig. 5a-b**, pLSM imaging produced high-resolution images of the entire preparation, facilitating systematic quantification. Comparative analysis across samples revealed significant heterogeneity in shape, cell number, and Sox2 expression profiles (**Fig. 5c**), underscoring the importance of such high-throughput approaches for generating statistically significant insights. Furthermore, we applied the same sample mounting and imaging strategy to perform two-color high-resolution imaging of human iPSC-derived vessel organoids labelled with CD31 antibody (an epithelial cell marker) and ZO1-mEGFP expression (a tight junction protein reporter^27^). As shown in **Fig. 5d** and **Supplementary Video 6**, this approach unveiled sub-cellular resolution morphological details across multiple vessel organoid samples. Additionally, a comparison of CD31 staining and ZO1-mEGFP expression pattern revealed lower levels of ZO1-mEGFP signal in the peripheral sprouting vessels, consistent with their lack of tight gap junctions^28^.

Overall, the results from these high-throughput experiments, combined with rigorous quantification, demonstrated the scalability and effectiveness of the pLSM framework for enabling quantitative high-throughput phenotyping of different types of organoid samples at sub-cellular resolution. Through its ability to capture a comprehensive view of the molecular and cellular heterogeneity within these complex 3D structures, pLSM has the potential to facilitate crucial insights into the intricate mechanisms that underlie various diseases.

### Unraveling the cellular dynamics across bacterial pellicle biofilms at the air-liquid interface

By utilizing the light sheet thickness modulation and live imaging capabilities of pLSM, we aimed to gain insights into the organization of bacterial biofilms and the underlying cellular dynamics. Biofilms represent the most prevalent form of multicellular organization in the bacterial world, with important implications for biotechnology applications (e.g., waste degradation and chemicals production), clinical relevance, and bio-geochemical cycling in water and soil environment ecosystems^29, 30^. The intricate interactions (e.g., social and electrical connections) among bacterial cells within biofilms result in emergent properties that extend beyond the sum of individual components^30^. Therefore, understanding the complex cellular dynamics underlying biofilm formation, maintenance and function is crucial.

Recent live imaging studies on bacterial biofilms^29^ have provided crucial insights into their dynamic nature, such as the remarkable transformations from 2D branched morphology to densely packed 3D clusters^31^, as well as the intricate cellular dynamics underlying biofilm formation^32^. However, most of these investigations have been restricted to imaging through the biofilm surface, resulting in lower axial resolution and limited imaging depth. As a result, our understanding of the cellular dynamics throughout the thickness of biofilms remains limited. Furthermore, the ability to analyze the effects of various mutations within biofilms would greatly benefit from the capacity to map changes across different layers.

To address these limitations, we developed and optimized a novel imaging assay that enables live imaging of thick cross-sections through pellicle biofilms formed at the air-liquid interface by *Pseudomonas aeruginosa*^33^ (**Fig. 6, Supplementary Videos 7-9**). By cultivating the pellicles in glass cuvettes (**Fig. 6a**), we achieved optical access from multiple sides. Leveraging the software-driven light sheet thickness modulation capability of pLSM, we fine-tuned the planar illumination thickness to capture the dynamic cellular processes within a thick optical section. We performed live imaging of pellicles derived from 2.5% fluorescently labeled (mScarlet) *Pseudomonas aeruginosa* cell pools^33^. To minimize the light-induced toxicity while still capturing the fast cellular dynamics over large field-of-view, we carefully tested a range of imaging speeds, resulting in a 2 Hz sampling rate. This enabled extended imaging sessions while facilitating cell tracking over time (**Supplementary Videos 7-8**). Moreover, we conducted long-term recordings lasting up to 7 hours, capturing images at 12-second intervals (**Supplementary Video 9**). The resulting large datasets were analyzed by using Ilastik^34^ framework for pixel classifications into three categories: tight clusters at air-liquid interface, individual cells or small cluster, and background regions (**Fig. 6a**).

**Figure 6.**
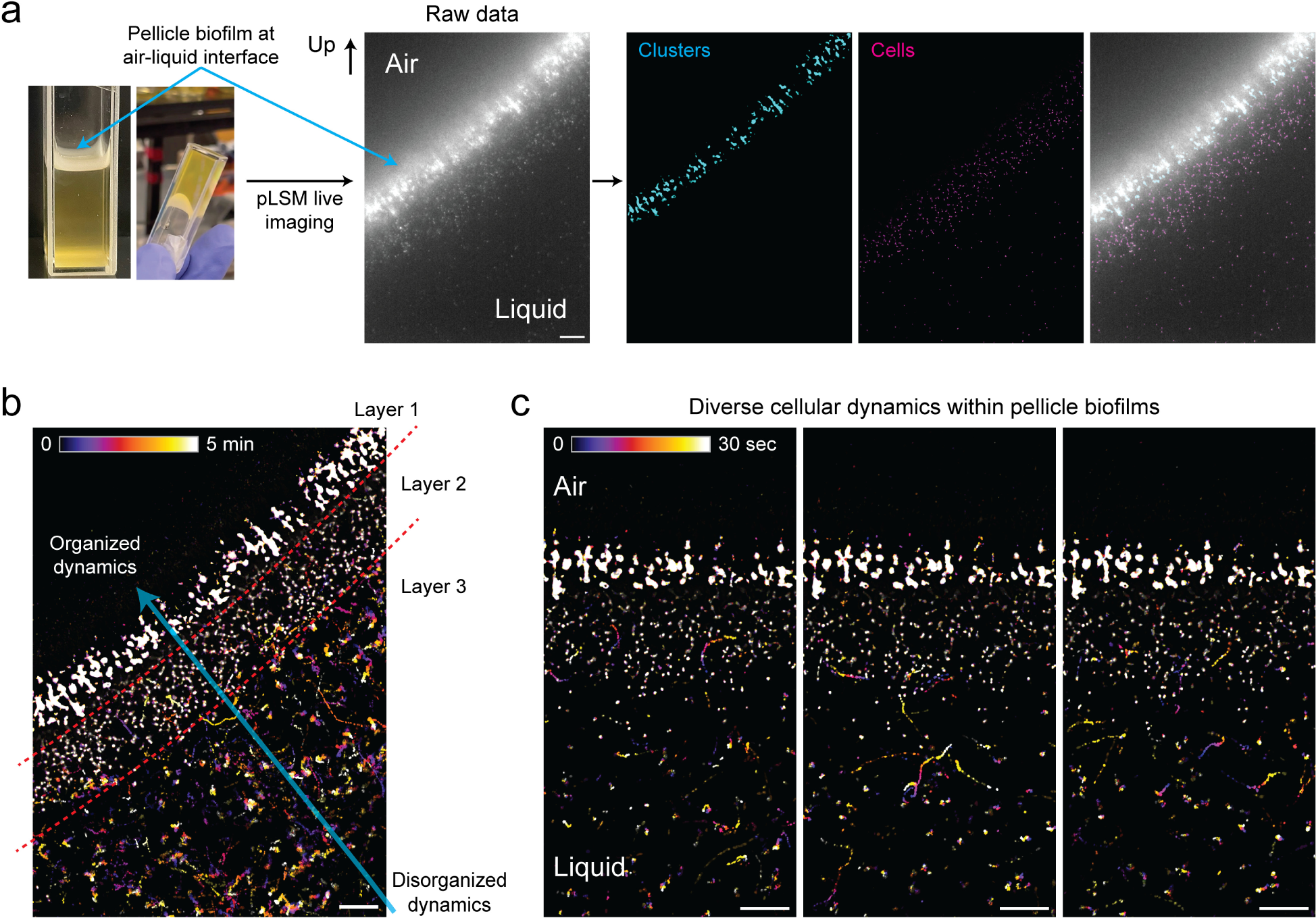
Mapping the dynamic layered architecture of bacterial pellicle biofilms at the air-liquid interface. (a) *Pseudomonas aeruginosa* pellicle biofilms were generated in cuvettes for live imaging of a thick optical cross-section through the depth. Images show representative raw data and detection of clusters and cells. (b) Color-coded projections across time to visualize the dynamic layered architecture of biofilm pellicles. Three distinct dynamic layers are apparent, showing disorganized to organized dynamics. (c) Color-code projections across time, at three representative time points, showing examples of cellular dynamics. Also see **Supplementary videos 7-9**. All scale bars are 50 µm.

Our findings revealed the presence of three morphologically and dynamically distinct regions within *Pseudomonas aeruginosa* pellicles (**Fig. 6b-c**). Layer 1 comprised tightly clustered cells located at the air-liquid interface, exhibiting highly dynamic shape changes. Layer 2 exhibited a heterogeneous porous organization characterized by chain-like arrangements of bacterial cells, allowing rapid cellular movements in specific regions while impeding motility in others. Layer 3 encompassed freely floating bacteria within the liquid, displaying varying degrees of motility, ranging from near immobility to high-speed projectile-like motions. Notably, we observed significant cell exchange between layers, particularly between layers 2 and 3, indicating intricate cellular interactions within the biofilm as a single system.

In summary, our comprehensive live imaging experiments demonstrate the unique compatibility of pLSM in mapping the intricate cellular dynamics and organization present throughout the depth of bacterial biofilms. These findings contribute to a deeper understanding of biofilm biology, with significant implications in clinical, geo-biological, and biotechnology contexts.

## DISCUSSION

We have introduced a novel LSFM imaging framework, called pLSM, which offers a highly cost-effective and scalable solution for imaging of living and large cleared biological samples. By leveraging off-the-shelf commercial-grade components, we have heavily reduced the implementation cost while ensuring high imaging quality, making pLSM a cost-effective alternative to high-end LSFM systems. We demonstrated the robustness and versatility of pLSM by multi-color imaging and quantitative analysis of diverse types of samples, including intact mouse brains, human brain tissue, brain and vessel organoids and bacterial biofilms. Additionally, we demonstrated the compatibility of pLSM with different detection objectives (e.g., for higher NA imaging).

The pLSM framework also incorporates several unique advanced features. Firstly, the software-driven modulation of the light sheet thickness allows for easy adjustment of axial resolution to fit the experimental need (e.g., for iterative multi-scale imaging of very large samples). This feature proved particularly effective for gentle live imaging of the dynamic cellular processes within bacterial pellicle biofilms formed at the air-liquid interface, providing crucial insights into their layered architecture. Secondly, the implementation of a generic over-the-network control enables remote operation and, together with compact hardware footprint, facilitates easy adoption in new locations. Thirdly, the linearly adaptive light sheet offset correction approach, by utilizing only a few pre-defined points, optimizes the sample imaging time and significantly advances the offset calibration strategy previously introduced in COLM system^2, 3^. Importantly, this approach also ensures compatibility with various imaging scenarios, encompassing different optical properties such as water-based, iDISCO, and CLARITY techniques.

However, there are certain limitations to consider. The imaging speed in pLSM is partly constrained by the refresh rate of the projectors used. In our implementation, we utilized projectors with a 60 Hz refresh rate, which generally suffices for most applications. Nevertheless, it is worth noting that scanning speeds achieved in pLSM may not reach the levels attainable with fast resonant galvo scanners. Future enhancements could involve integrating more advanced projectors to further boost the imaging speed needed for specific experimentations. Similarly, while pLSM is compatible with the commonly fluorophores in biological sciences (e.g., we imaged GFP, eYFP, mScarlet, and Alexa647 dyes), the excitation wavelengths may not be ideal for all flurophores of interest. Additionally, the imaging is currently limited to three colors, but future improvements may involve customization of the laser diode sets in pocket projectors.

In conclusion, pLSM provides an easily scalable, cost-effective, and versatile high-resolution imaging framework. Its novel features, such as ease of light sheet thickness modulation, fast adaptive offset corrections, and over-the-network control architecture, make it uniquely suitable for several experimental needs. Beyond accessibility and versatility, pLSM opens new possibilities for high throughput imaging applications, including drug screening and characterization of a large number of diseased brains and organoids.

## METHODS

### pLSM implementation

#### pLSM setup

The pLSM system consists of two illumination and one detection arms (**Fig. 1**), mounted on 60 mm cage assembly system (Thorlabs) (**Supplementary Table 1** lists all components). Each illumination arm utilizes a MEMS LASER projector (e.g., Nebra Anybeam), mounted on two dovetail translation stages (DT12) for precise adjustment along the light path. The projector incorporates TO-8 LASER diodes that provide blue (440-460nm), green (515-530nm), and red (632-642nm) colors. A combination of two achromatic doublets (AC254-045-A) forms a Plössl-style scan lens with a focal length of approximately 22.5mm. An achromat doublet (AC508-075-A) was paired with an air objective (Mitutoyo 10x Plan Apo) for illumination. In the detection arm, the Mitutoyo 10x Plan Apo or ASI 16.67x/0.4 NA objective is used, along with a tube lens (TTL200) and a CMOS camera (GS3-U3-89S6M-C, FLIR) with a pixel dimension of 3.45 µm. A multi-band filter (577/690nm, #87-251, Emund Optics or 432/515/595/730 nm, Semrock) was used as the emission filter. To match refractive index, a custom-designed oil chamber is 3D printed using polylactic acid (PLA) and features windows for illumination and detection, sealed with cover glass (VWR). A refractive index matching oil with an RI of 1.454 (Cargille) is used to fill the chamber. Sample positioning is achieved using a motorized stage (PT3 Z8, Thorlabs) for fine adjustments during imaging, along with additional 3-degree-of-freedom dovetail translation stages (DT12XYZ, Thorlabs) and a rotation stage (QRP02, Thorlabs) for additional flexibility during sample mounting. The sample is mounted in a quartz glass cuvette (FireflySci). The imaging workflow is implemented on Nvidia Jetson Nano board, which provides four high-speed USB 3.0, one HDMI, one DP, and one Ethernet ports. The HDMI ports of the two projectors are connected to the board’s HDMI and DP display ports using an HDMI-HDMI cable and an active DP to HDMI cable, respectively. The projectors are powered through USB using Micro USB cables connected to two USB ports on the board. The camera is connected to the board’s third USB port via a USB Micro B to USB Type A cable. A solid state hard drive is connected to the fourth USB port using a USB Type-C to USB cable. The board is connected to the control computer via an Ethernet cable for remote control. The motorized stages are connected to motor control boxes (KDC 101), which are then connected to the control computer.

#### pLSM control

The control software, implemented in Python, comprises several classes that handle different aspects of the system: (a) the Illumination Class controls the light sheet using the Xlib protocol to adjust the projector output, enabling precise control over the illumination settings; (b) the Camera Class utilizes the Spinnaker package to control the FLIR camera, allowing for parameter configuration and imaging trigger control; (c) the Widget Class is responsible for creating the software’s graphical user interface (GUI) using the IPyWidgets package, providing an interactive control interface for the user; (c) the Experiment Class is responsible for setting all the experiment parameters, executing actions, and saving acquired images directly to the hard drive via USB 3.0. It coordinates the entire imaging process. Remote control of the software on the Nvidia Jetson Nano board is achieved through SSH protocol for secure remote access. The control GUI can be accessed through a web browser, providing convenience and flexibility in controlling the pLSM system remotely. For precise control of the motorized stage, the thorlabs_apt Python package was utilized. The motorized stage’s movement is synchronized with the camera acquisition through the Experiment Class, ensuring precise alignment and coordination during the imaging process.

#### System PSF characterization

The point spread function of the system was measured by imaging micro-meter sized fluorescent beads (17154-10, PolyScience), prepared by a 1:1million dilution in 0.8% low melting agarose solidified in quartz cuvette. The point spread function was imaged by moving the beads in axial direction at 1µm step size. Different number of light sheet lines were used to test the tunable light sheet profiles. For the beads data analysis, images were first processed by ilastik^34^ to generate beads mask, which were then used to determine the center of mass coordinates for each bead. Beads from the original image were cropped and aligned based on the center of mass, followed by full width half maximum calculations.

#### Image processing

Median filter with radius 0.1 was applied to remove hot pixel noise. The image tiles were stitched using terastitcher^35^ or bigstitcher^36^. The volume rendering was performed with Amira. For TH segmentation, suiteWB pipeline^22^ was used as described previously^22^. Briefly, the whole brain datasets were registered onto a local average reference atlas. The registered data was split into dense and sparse regions for multi-model segmentation. Finally, the detected cells were annotated with Allen brain atlas (ABA, ccfv3) regions for further comparison.

### Fixed brain sample preparation

#### Brain samples

For mouse dopamine and vasculature labeling, CD1 WT (Crl:CD1(ICR), charles river #022) mice were used. *Thy1-eYFP* (B6.Cg-Tg(Thy1-YFP)HJrs/J, JAX Strain 003782) mice were used for the CLARITY sample. All experiments were performed with 8-10 weeks old mice. For the human vasculature samples, post-mortem frozen de-identified human cortex sample was used. All experimental procedures were approved by the IACUC at Columbia University.

#### Tissue fixation

The mouse brains were perfused with 4% PFA and extracted, followed by overnight fixation in 4% PFA at 4℃. Freshly frozen human brain samples were incubated in 4% PFA for one day at 4℃. All samples were washed and stored in 1xPBS at 4℃.

#### iDISCO sample preparation

Standard iDISCO^19^ procedure was used, consisting of sample pretreatment, immunolabeling and clearing steps. For pretreatment, brain samples were dehydrated in 20%, 40%, 60%, 80%, 100%, 100% v/v methanal/H_2_O series for 1h each, followed by 66% v/v dichloromethane (DCM)/methanal incubation overnight. The samples were then washed by 100% methanal 1h twice and incubated in 5% H_2_O_2_ v/v in deionized water overnight at 4℃. Then rehydration was performed in 80%, 60%, 40%, 20% v/v methanal/H_2_O and PBS for 1h each. Before immunolabeling, the brain samples were first washed in PBS+0.2% Triton 1h for twice, followed by permeabilization in PBS+0.2% Triton at 37℃ for 1 day. The samples were then blocked in 1% BSA in PBS for 1 day at 37℃. Antibody labelings were performed in PBS with 0.1% Triton / 1% BSA and corresponding primary antibodies. Anti-TH antibody (ab113, Abcam) was used at 1:500 dilution. The mouse brain vasculature labeling was performed similar to previously reported procedure^21^, by using a cocktail of antibodies: goat anti-podocalyxin (1:1500, AF1556, R&D Systems), goat anti-CD31 (1:300, AF3628, R&D Systems) and rabbit anti-Acta2 (1:1000, ab21027, Abcam). Human vasculature labeling was performed by using a cocktail of goat anti-podocalyxin (1:20, AF1658, R&D Systems), mouse anti-CD31 (1:50, BBA7, R&D Systems) and rabbit anti-Acta2 (1:1000, ab21027, Abcam). The brain samples were incubated in primary labeling solution for 10 days at 37℃, with one solution replacement at the end of day 5. The brains were washed in PBS + 0.1% Triton for 5 times at least 1h each and until the next day. For the mouse TH labeling, donkey anti-sheep Alexa 647 (1:1000, A-21448, Thermofisher) was used; for mouse vasculature labeling, donkey anti-goat Alexa 647 (1:500, A-21447, Thermofisher) and donkey anti-rabbit Alexa 647 (1:500, A-32795, Thermofisher) were used. For human vasculature labeling, donkey anti-goat Alexa 647 (1:500, A-21447, Thermofisher), donkey anti-mouse Alexa 647 (1:500, A-32787, Thermofisher) and donkey anti-rabbit Alexa 555 (1:500, A-31572, Thermofisher) were used. Secondary antibody incubation was also done for 10 days at 37℃, with one solution replacement at the end of day 5. For clearing, samples were dehydrated in 20%, 40%, 60%, 80%, 100%, 100% v/v methanal/H_2_O for 1h each step followed by incubation in 66% v/v DCM/methanal for 3h. The samples were then washed in DCM 15min twice and put in dibenzyl ether (DBE, 33630, Sigma-Aldrich, RI=1.562) for refractive index matching until the next day for imaging.

#### Thy1-eYFP CLARITY sample preparation

PFA fixed brain samples were incubated in 1% hydrogel monomers (HMs) consisting of 1% w/v acrylamide (#1610140, BioRad), 0.05% w/v bisacrylamide (#1610142, BioRad), 4% paraformaldehyde (15710-S, Electron Microscopy Sciences), 1xPBS, deionized water, and 0.25% of thermal initiator (VA-044, Fisher Scientific) at 4℃ overnight followed by oxygen removal and replacement of nitrogen using vacuum pump. The air-tight sample was then polymerized at 37℃ for 6 hours. Then the sample was undergone passively clearing by incubating in SBC buffer at 37℃ with gentle shaking. Sample transparency was assessed to determine clearing completion, which usually took about 4-6 weeks. Sample was washed in 0.2M boric acid and can be left in upon imaging. RapiClear (SunJin lab, RI=1.47) was used for RI matching before imaging.

### Bacterial strains and pellicle preparation

*Pseudomonas aeruginosa* strains used in this study are UCBPP-PA14^37^ (WT, LD0) and PA14 P_PA1/04/03_-mScarlet^33^ (LD4764). Liquid cultures were grown in Lysogeny Broth (LB) at 37 °C with shaking at 250 rpm. Overnight precultures were diluted 1:100 in LB and grown to mid-exponential phase (OD at 500 nm ∼ 0.5). OD values at 500 nm values were read in a Spectronic 20D+ spectrophotometer (Thermo Fisher Scientific [Waltham, MA]) and cultures were adjusted to the same OD. Adjusted cultures were then mixed in a 2.5:97.5 ratio of fluorescent:non-fluorescent cells. Two mL of mixed culture was added to a 10 mm optical glass Type 1FL Macro Fluorescence Cuvette (1FLG10, Fireflysci). The cuvette was covered with parafilm and grown at 25°C for 3 days. The live imaging was performed with pLSM using ASI 16.67X/0.4 NA objective and 4 pixels thick light sheet. The raw data was segmented by Ilastik^34^ framework by classifying pixels into tight clusters at air-liquid interface, individual cells or small cluster, and background regions. The color-coded projections across time were generated by using ImageJ/Fiji.

### Brain and vessel organoids

#### Culture of hiPSCs

The hiPSC line Edi042A was purchased from Cedars Sinai. The hiPSC line WTC-ZO1-mEGFP (AICS-0023)^27^ was a gift from the Gordana Vunjak-Novakovic lab. Both cell lines were maintained in six-well plates coated with Matrigel growth factor reduced basement membrane matrix (GFR-Matrigel, Corning, 354230) in mTeSR plus medium (STEMCELL Technologies, 100-0276). To coat the six-well plates, 1.5 mL of GFR-Matrigel (diluted at a 1:100 ratio with DMEM/F12 medium, Gibco, 11320033) was added per well and incubated at 37°C for 1h. To passage hiPSCs (60-70% confluency), cells were rinsed with 3-4 mL of DMEM/F12 medium per well, and then 2 mL of Accutase (Sigma-Aldrich, A6964) was added for 5-6 min at 37°C. Subsequently, 2 ml of mTeSR plus medium was added to neutralize the dissociation. Cells were collected after centrifugation at 1000 rpm for 3 min. Cells were resuspended with mTeSR plus with 10 μM Y27632 (Selleckchem, S1049) and evenly distributed on GFR-Matrigel coated wells.

#### Generation of brain organoids

The brain organoids were generated from the Edi042 hiPSC line based on a previously described method with minor modifications^38, 39^. On day 0, hiPSCs were incubated with Accutase at 37°C for 7 min and dissociated into single cells. Approximately 10,000 single cells were added to a 96-well U-shaped-bottom low-attachment plate per well (Thermo Scientific, 174925) in mTeSR plus medium with 10 μM Y27632, and incubated at 37°C with 5% CO_2_. From day 1 to day 6, the medium was replaced with neural induction medium (NIM) daily, containing DMEM/F12 medium, 20% Knockout Serum Replacement (KSR, Gibco, 10828028), 1% Minimum Essential Medium non-essential amino acids (MEM-NEAA, Gibco, 11140050), 0.1 mM 2-mercaptoethanol (Gibco, 21985023), 50 μg/mL Gentamycin (Gibco, 15750060), 10 μM SB431542 (Selleckchem, S1067) and 5 μM dorsomorphin (Medchemexpress, HY-13418). From day 7 to day 26, the medium was replaced with neural differentiation medium every other day, containing neurobasal medium (Gibco, 21103049), 2% B27 supplement minus vitamin A (Gibco, 12587010), 1% GlutaMAX (Gibco, 35050079), 50 μg/mL Gentamycin, 20 ng/mL EGF (ProSpec, CYT-217) and 20 ng/mL FGF2 (ProSpec, CYT-557). From day 27, the EGF and FGF2 were replaced with 20 ng/mL of brain-derived neurotrophic factor (BDNF, ProSpec, CYT-1081) and 20 ng/mL of neurotrophin-3 (NT3, ProSpec, CYT-257).

#### Generation of vessel organoids

The vessel organoids were generated from the WTC-ZO1-mEGFP line as previously described with minor modifications^24, 40^. On day 0, hiPSCs were dissociated into single cells with Accutase. The collected cells were resuspended in mTesR plus medium with 10 μM Y27632. 1,000 cells were reaggregated in a 96-well U-shaped-bottom low-attachment plate per well. On day 1, the medium was replaced with N2B27 medium, containing 48.5% (v/v) DMEM/F12 medium, 48.5% (v/v) neurobasal medium, 0.5% N2 supplement (Gibco, 17502048), 1% B27 supplement minus vitamin A, 0.5% MEM-NEAA, 0.5% GlutaMAX, 0.1 mM 2-mercaptoethanol, 50 ug/mL Gentamycin, 12 μM CHIR99021 (Tocris, 4423) and 30 ng/mL BMP4 (ProSpec, CYT-1093). On day 4, the medium was replaced with N2B27 medium + 100 ng/ml VEGFA (Peprotech, 100-20) + 2 μM forskolin. On day 6, the medium was replenished with N2B27 medium + 100 ng/ml VEGFA + 100 ng/ml FGF2 every other day.

#### Immunostaining

Brain organoids at day 37 and vessel organoids at day 14 were fixed with 4% paraformaldehyde (Electron Microscopy Sciences, 15713) at 4°C overnight. The organoids were washed three times with phosphate-buffered saline (PBS, Corning 46013CM), blocked in 10% donkey serum for 2h and incubated with primary antibodies at 4°C overnight. After aspirating the primary antibodies, the organoids were washed with PBS Tween-20 buffer (PBST, Thermo Fisher, 28352) three times for 30 min and incubated with secondary antibodies at 4°C overnight. The organoids were washed three times with Tris-buffered saline Tween-20, (TBST, Thermo Fisher, 28360) and stained with 4′,6-Diamidino-2-phenylindole dihydrochloride, 2-(4-Amidinophenyl)-6-indolecarbamidine dihydrochloride (DAPI, Millipore-Sigma, D9542) for 15 min.

#### Tissue clearing

Organoids were embedded in 1% agarose gel and lined up before tissue clearing. The stained organoids were cleared by FDISCO as previously described^26^. Samples were dehydrated with tetrahydrofuran (Sigma, 676764) solutions at graded concentrations: 50% (v/v), 70% (v/v), 80% (v/v), 100% (v/v) and 100% (v/v), with 1h for each step and incubated at 4°C. Then the samples were transferred to dibenzyl ether (DBE, Sigma, 33630) and incubated for 1h at room temperature.

#### Antibodies

The antibodies used in this study are listed as follows with the dilution ratios: goat-anti-human SOX2 (R&D, AF2018, 1:200), rabbit-anti-human CD31 (Abcam, ab28364, 1:200); Alexa Fluor 647 Donkey anti-goat IgG (Jackson Immnoresearch Lab, 705-606-147, 1:1000), Alexa Fluor 488 Donkey anti-rabbit IgG (Jackson Immnoresearch Lab, 711-546-152, 1:1000), Alexa Fluor 647 Donkey anti-rabbit IgG (Thermo Fisher, A32795, 1:1000).

## Acknowledgements

We are grateful to Roberto Etchenique for initial discussions related to LASER projectors, and thankful to Maura Boldrini for providing fixed human brain tissue sample. This work was supported by NIH grants DP2MH119423 and UH3TR002151 to R.T.; UH3TR002151 and UH3NS115598 to K.W.L.; R01AI103369 to L.E.P.D.; and by Columbia University Arts & Sciences startup grant to R.T..

## Author Contributions

Y.C. and R.T. conceptualized and designed the pLSM framework. Y.C. implemented and characterized pLSM, with inputs from C.G. on optical simulation and CAD design, and from S.C. on initial prototyping. The imaging experiments were conducted as follows: Y.C., S.C., E.D.D., C.G., M.S.D., and R.T. prepared mouse and human brain samples, and performed pLSM imaging. C.X., S.C., R.T., and K.W.L. generated and imaged the brain and vessel organoids. S.C., H.D., L.E.P.D., and R.T. developed the live imaging assay for pellicle biofilms and performed the live imaging experiments. R.T. and Y.C. analyzed all the data and wrote the paper, with input from all the authors. R.T. supervised the project.

## Data availability statement

All datasets will be made available on request.

## Code availability

All the code is made available publicly as a GitHub repository: https://github.com/tomerlab/pLSM-Control

## Completing interests

Authors declare no competing interests.

