## Supplementary Table and Video Legends for "Scalable projected Light Sheet Microscopy for high-resolution imaging of living and cleared samples"

**Supplementary Table 1. Breakdown of pLSM cost.** pLSM achieves one twentieth to one fourth of the cost compared to other systems. For reference, COLM system costs > $250k, dual color dual illumination OpenSPIM is ~$50k (https://www.bioimager.com/product/biopenspim-light-sheet-microscope) and mesoSPIM can cost $170k to $240k (https://mesospim.org/faq/#how-much-does-a-mesospim-cost).

| Part | Item | Usage | Count | Price $ | Total $ |
|---|---|---|---|---|---|
| **Illumination** | AnyBeam Projector | Laser source | 2 | 200 | 400 |
| | Dovetail Stage DT12 | Projector mounting | 4 | 84 | 336 |
| | AC254-045-A | Plössl scan lens | 4 | 162 | 648 |
| | AC508-075-A | Tube lens | 2 | 129 | 258 |
| | Mitutoyo 10x Plan Apo | Objective | 2 | 880 | 1760 |
| **Detection** | TTL200 | Tube lens | 1 | 500 | 500 |
| | GS3-U3-89S6M-C | Camera | 1 | 3095 | 3095 |
| | Mitutoyo 10x Plan Apo (Optional: ASI 16.6X/0.4NA) | Objective | 1 | 880 | 880 |
| **Other** | Nvidia Jetson Nano | System | 1 | 300 | 300 |
| | Dovetail Stage DT12 | Sample mount | 3 | 84 | 252 |
| | Rotation Stage QRP02 | Sample mount | 1 | 160 | 160 |
| | Tubes, rails, bars | Structure | 1 | 200 | 200 |
| | KDC 101 (optional) | Motor control | 3 | 758 | 2274 |
| | PT3 Z8 (optional) | Motor stage | 1 | 3030 | 3030 |
| **Total** | | | | | 8,809-14,113 |

**Supplementary Videos:**

**Supplementary Video 1. Whole-brain volumetric rendering of TH+ neurons mapped using projected Light Sheet Microscopy (pLSM)**. This representative example showcases the brain-wide mapping of TH+ neurons in an iDISCO+ cleared sample.

**Supplementary Video 2. Visualization of whole-brain segmentation of TH+ neurons in the same sample imaged with both pLSM and COLM systems.** This representative example demonstrates a high degree of agreement in the automated segmentation of data acquired from both pLSM and COLM systems from the same sample.

**Supplementary Video 3. Whole-brain rendering of *Thy1-eYFP* transgenic mouse brain imaging using pLSM with CLARITY clearing.** This representative example showcases high-resolution imaging of CLARITY-cleared *Thy1-eYFP* transgenic mouse brain images using pLSM.

**Supplementary Video 4. Volume rendering of *Thy1-eYFP* transgenic mouse brain imaging using pLSM with CLARITY clearing.** This representative small volume rendering showcases the high-resolution imaging using pLSM.

**Supplementary Video 5. Volumetric rendering of the pLSM imaging of an ensemble of human iPSC-derived brain organoids.** The video shows mounting and pLSM imaging of an ensemble of 8 brain organoid samples, immunostained with anti-Sox2 antibody, in a single imaging session.

**Supplementary Video 6. Volumetric rendering of human iPSC-derived vessel organoids imaged with pLSM.** The video shows pLSM imaging of vessel organoids stained with anti-CD31 antibody.

**Supplementary Video 7. Live imaging of bacterial biofilm pellicle at the air-liquid interface.** Representative movie showing cellular dynamics across the *Pseudomonas aeruginosa* pellicle biofilm at the air-liquid interface.

**Supplementary Video 8. Live imaging of bacterial biofilm pellicle at the air-liquid interface.** Representative movie, at different location then Supplementary Video 7, showing cellular dynamics across the *Pseudomonas aeruginosa* pellicle biofilm at the air-liquid interface.

**Supplementary Video 9. Live imaging of a highly motile bacterial biofilm pellicle at the air-liquid interface.** Representative movie of a highly motile pellicle biofilm formed by *Pseudomonas aeruginosa* at the air-liquid interface.
